# Temperature and dietary energy content influence female maturation age and egg nutritional content in Atlantic salmon

**DOI:** 10.1101/2022.09.09.507230

**Authors:** Katja S. Maamela, Eirik R. Åsheim, Paul V. Debes, Andrew H. House, Jaakko Erkinaro, Petra Liljeström, Craig R. Primmer, Kenyon B. Mobley

## Abstract

The environment experienced by a female influences reproductive traits in many species of fish. Environmental factors such as temperature and diet are not only important mediators of female maturation and reproduction but also of egg traits and offspring fitness through maternal provisioning. In this study, we use three-year-old, tank-reared, Atlantic salmon from two Finnish populations to investigate the effect of temperature and diet on maturation and egg traits. We show that a temperature difference of 2°C is sufficient to delay maturation in female Atlantic salmon whereas a 22% reduction in dietary energy content had no effect on maturation. Diet did not influence the body size, condition, or fecundity of the mature females or the size or protein content of the eggs. However, a higher energy diet increased egg lipid content. Neither female body size nor condition were associated with egg size or fat/protein composition. Our results indicate that female salmon that have a poorer diet in terms of energy content may have a reproductive disadvantage due to lower energy provisioning of eggs. This disadvantage has the potential to translate into fitness consequences for their offspring.

## Introduction

The environment a female experiences prior to and during maturation is key in mediating reproductive traits across fish species (Green, 2008). Ecological factors, such as temperature and resource acquisition, directly affect female body size and condition both of which are tightly linked with sexual maturation and reproductive output in many species (Gregersen *et al*., 2011; Kjesbu *et al*., 1998; Sink & Lochmann, 2008; Yoneda & Wright, 2005). As energy needed for reproduction is derived from maternal resources (Wootton & Smith, 2015), different environmental factors such as diet may influence egg size, maternal provisioning of the eggs, and subsequent offspring growth and survival (Gagliano & McCormick, 2007; Mazorra *et al*., 2003; Sink & Lochmann, 2008). Therefore, the environment a female inhabits not only affects her reproductive fitness but also the fitness of her offspring through maternal provisioning.

Atlantic salmon, *Salmo salar*, provide an excellent model system to investigate factors affecting sexual maturation and reproductive traits in females (reviewed in Mobley *et al*., 2021). The age at which a salmon reproduces for the first time, i.e., age at maturity, is influenced by a number of environmental factors including water temperature, photoperiod, and diet (Fjelldal *et al*., 2011; Imsland *et al*., 2014; Jonsson & Jonsson, 2011; Taranger *et al*., 2010), as well as by the underlying genetic architecture (Ayllon *et al*., 2015; Barson *et al*., 2015; reviewed in Mobley *et al*., 2021; Sinclair-Waters *et al*., 2020). Atlantic salmon have an increased growth rate, lipid accumulation, and body condition in warmer water temperatures which is generally associated with lower maturation age (Duston & Saunders, 1999; Jonsson *et al*., 2013, 2012; Jonsson & Jonsson, 2011). As the maturation process is connected with the fish’s growth and ability to utilize its somatic energy reserves (Jonsson *et al*., 2013) and the energy stores need to be sufficient for maturation to occur (Rowe *et al*., 1991), the effect temperature has on maturation age may additionally be linked with resource acquisition through diet and feeding opportunities. The acquired somatic energy stores are necessary for the development of both primary (ovaries) and secondary (morphological and colour changes) sexual characters as well as for return migration and activities associated with spawning, such as digging of nests for eggs in the river bed (called ‘redds’), deposition of eggs within the redds, and territorial behaviour especially towards other females (Fleming, 1996; Jonsson & Jonsson, 2011; Taranger *et al*., 2010; Thorpe *et al*., 1998).

In Atlantic salmon females, ovarian maturation typically takes place over a four-month period prior to spawning (Jonsson & Jonsson, 2011). During this period, the oocyte is supplied with the necessary components needed for the early development and growth of the offspring and the mature oocyte is ovulated into the body cavity (Wootton & Smith, 2015). Adults stop feeding during the return migration (Jonsson *et al*., 1991; Kadri *et al*., 1996) and females utilize their somatic nutritional reserves acquired prior to migration to provision eggs (Brooks *et al*., 1997). As adequate lipid stores are needed for the energy-intensive processes of maturation and reproduction, and energy allocated to gonads is approximately 30% of the total somatic energy in female salmon, a high-lipid diet may be associated with a higher maturation rate in females (Jonsson *et al*., 2013; Jonsson & Jonsson, 2003).

Diet may additionally influence female reproductive fitness via fecundity, i.e., the number of eggs a female can produce, (Bagenal, 1969; Eskelinen, 1989; Izquierdo *et al*., 2001; Scott, 1962) and egg size (Sink & Lochmann, 2008). A high-lipid diet promotes both rapid growth and lipid accumulation (Jonsson *et al*., 2012; Kadri *et al*., 1996; Shearer *et al*., 2006) as well as larger body size at maturation which may then lead to higher fecundity and egg size in females (Barneche *et al*., 2018; Fleming, 1996; Heinimaa & Heinimaa, 2004). Increased egg size, in turn, is associated with a higher chance of initial survival for the offspring as juveniles hatching from larger eggs also tend to be bigger (Einum, 2003; Einum & Fleming, 2000; Thorpe *et al*., 1984) and have an initial higher growth rate due to the eggs holding greater energy stores to be used during early life-stages (Brooks *et al*., 1997; Einum, 2003).

Maternal diet and acquired somatic energy resources likely influence egg quality through maternal provisioning of yolk in eggs (Bogevik *et al*., 2012; Brooks *et al*., 1997; Kamler, 1992; Kjørsvik *et al*., 1990). The yolk is the main element of fish eggs supporting the embryo’s growth and development in teleost fishes. The yolk is composed of various components such as lipids, proteins, and carbohydrates as well as micronutrients and maternal mRNA (Brooks *et al*., 1997; Kamler, 2008). Of the essential yolk components, lipids and proteins are found in the highest concentrations (Brooks *et al*., 1997). The yolk is not only a vital resource for the embryo during development, but also for the larvae after hatching as an endogenous energy source for growth and maintenance until the start of independent feeding in many species of fish (Berg *et al*., 2001; Kamler, 1992, 2008; Wootton & Smith, 2015). Atlantic salmon females exhibit no parental care beyond their choice of nest site (Jonsson & Jonsson, 2011) thus emphasizing the importance of maternal provisioning of the offspring.

Given the importance of energy resources during early development, maternal energy provisioning of the eggs influences both offspring fitness and maternal reproductive fitness. Currently, our knowledge of egg characteristics, specific nutritional requirements by the offspring, and the effect of diet on female maturation and egg quality in Atlantic salmon is limited to a handful of studies that have measured the egg energy composition (Berg *et al*., 2001; Einum & Fleming, 2000; Heinimaa & Heinimaa, 2004; Rennie *et al*., 2005; Srivastava & Brown, 1991). Of the abovementioned studies, only Rennie *et al*. (2005) investigated the effect of female diet on egg composition. The previous studies on egg composition highlight the importance of maternal energy provisioning for the survival of the offspring during early development where an increase in egg energy stores is associated with lower mortality during incubation and a fitness benefit after hatching (Berg *et al*., 2001; Heinimaa & Heinimaa, 2004). How maternal diet is reflected in egg energy composition, on the other hand, is less well-understood.

In this study, we investigate how the environment influences reproductive traits of female Atlantic salmon. Specifically, we address how dietary energy content and water temperature affect maturity rate of the female salmon and how female body size, condition, fecundity, and egg traits are affected by dietary energy content. We hypothesise that diet should affect maturation and predict that females fed a diet of higher energy content should have more resources to allocate towards reproduction and show a higher rate of maturation. Similarly, we expect higher water temperature to lead to increased maturation rate. We also predict that the lipid content of the eggs from females on the higher energy diet would be greater compared to females receiving lower energy feed.

## Materials and Methods

### Study fish

The parents of the study fish were first-generation hatchery broodstock stocked in the Finnish rivers Kymijoki and Oulujoki with the Kymijoki fish originating from the river Neva in north-western Russia and the Oulujoki fish originating from several northern Baltic salmon populations, subsequently stocked in the river Oulujoki. Both rivers drain to the Baltic Sea. In this study, these two populations will be referred to as the Neva and Oulu populations, respectively. Details of the populations, crossing design, and rearing are explained in Åsheim *et al*. (2022). Briefly, unrelated fish were crossed within each population to create 52 Oulu crosses (families) and 68 Neva crosses. Eggs were fertilized in October 2017 and incubated in vertical flowthrough incubators at the Viikki campus of the University of Helsinki as described in Debes *et al*. (2020). Alevins were transferred to experimental facilities at the Lammi Biological Station (LBS, 61°04′45′′N, 025°00′40′′E, Lammi, Finland) in March 2018, several weeks after hatching but prior to independent feeding. Individuals from each family were divided approximately evenly among twelve 270 cm diameter circular tanks (maximum water volume 4.6 m^3^) which were supplied with a continuous flow of UV-sterilized water from the nearby lake Pääjärvi.

A temperature treatment was created by warming the water of half of the tanks by ∼1°C from ambient water temperature (warm) and cooling the other half by ∼1°C (cold) using a heat-exchange system in the inflow system thus creating a 2°C target temperature difference with the average water temperature across the study period in the warm treatment being 5.39 ± 3.30°C and in the cold treatment being 7.41 ± 3.31°C (mean ± SD) (see supplementary material Fig. S1). Egg trait components of this study focus on the females of the warm-water treatment because no females in the cold treatment matured during the 2020–2021 spawning season (see results). The light-regime followed the local natural cycle at LBS. All fish were initially fed fish feed (Hercules Baltic blend, Raisioaqua, Raisio, Finland) of appropriate pellet size for their mass *ad libitum*. Starting in July 2019, half of the tanks received one of two feed treatments: 1) control feed (as above) or 2) low-fat feed. The low-fat feed was manufactured with reduced fat content which resulted in ∼22% reduction in total energy content (kJ/100g) compared to the control feed (feed content analyses of each food batch conducted by Synlab Oy, Karkkila, Finland). Temperature and feed treatments were combined to create four treatment combinations which were evenly divided among 12 tanks resulting in three tanks for each feed x temperature combination.

Each individual was marked with a 12 mm Passive Integrated Transponder (PIT) tag to allow for individual identification and a fin clip was taken for parentage and population assignment as done in Åsheim *et al*. (2022).

### Female traits

The female body size was measured at three time points during the winter 2020 – 2021 (Table S1, Fig. S2) when the salmon were around three years old. Each female was first anaesthetized with tricaine methanesulfonate (MS-222) at a concentration of 0.125 g L^-1^ buffered with an equal amount of sodium bicarbonate before their fork length (body length, tip of the rostrum to midpoint in fork, mm) and wet mass (g) was measured. In addition to body size measurements, Fulton’s condition factor (K = 100*(M/L^3^), where M is wet mass in g and L fork length in cm) (Heincke, 1908), was calculated for all mature females.

### Maturation

Maturation of females was assessed during each of the three aforementioned measurement sessions (Table S1). Maturation stage of each female was categorized based on external characteristics (Table S2). Prior to egg stripping, the body wet mass of the mature females was recorded.

### Egg stripping

Mature females were stripped by gently applying pressure to the abdomen. Stripped eggs were collected into a clean 5 L bucket and placed in a 4°C refrigerator until further processing. During the third sampling period, females that had already been stripped once during the spawning season (sampling periods 1 or 2; Table S1) but had new eggs were restripped. If the number of eggs was found to be higher when the female was stripped for the second time compared to the first time, this was taken as an indication that the female had not been stripped close to ovulation time initially. These females (*n* = 2) were recorded as mature but were excluded from the final female and egg trait analyses.

### Egg weight and female fecundity

After stripping, ovarian fluid was removed by straining through a stainless-steel sieve (mesh size 1×2 mm) and remaining eggs were weighed to determine the total egg wet mass (g). The fecundity of each female was calculated by dividing the total egg wet mass by the wet weight of a single egg. The wet weight of a single egg was calculated as a mean based on the average weight (to 0.001 g accuracy) of five replicates of 10 eggs. Dry weight of the eggs was obtained by drying a sample of 10 eggs of known wet mass for 48 hours at 60°C until no change was observed in the sample weight. After drying, the sample was weighed to the nearest 0.0001 g and the mean individual egg weight was calculated from the total weight of the sample.

### Egg diameter

To measure egg diameter, 10 unfertilized eggs were placed on a petri dish on top of a continuous light source. The petri dish was covered with a light-blocking black box and the eggs photographed using a digital camera (Canon EOS 750D fitted with an EF-S 18-55 mm lens, aperture f/11, shutter speed 1/250s) with a mm ruler for scale. The egg diameter was calculated using *ImageJ* version 1.53s (Rueden *et al*., 2017) by calculating the mean diameter of the egg from eight measurements taken across each egg. The average egg diameter for each female was then calculated as the mean of the 10 eggs measured.

### Egg lipid content

Egg lipid content was quantified based on the lipid extraction method outlined in Shinn & Proctor, (2013). In brief, a sample of 10 frozen eggs from each female was homogenized in 900 µL of milliQ water using an Omni Bead Ruptor Elite (Omni International) after which the homogenate was transferred to a 15 mL glass vial and the residual was rinsed twice with 500 µL of ethanol. Lipids were extracted by adding 5 mL of hexane:isopropanol solution (3:2) to the homogenized eggs and immediately centrifuged at 6 238 g for 10 minutes. The top layer containing the extracted lipids in the hexane:isopropanol was pipetted to a new, pre-weighed 15 mL glass vial. The liquid phase was evaporated using nitrogen gas (Stuart SBHCONC/1 Sample Concentrator, Cole-Parmer, Staffordshire, UK) and the lipid residue (mg * wet weight^-1^) weighed to the nearest 0.0001 g.

### Egg protein content

Egg protein content was quantified using the Bradford method (Bradford, 1976) following the instructions of the Pierce(tm) Coomassie (Bradford) Protein Assay Kit (ThermoScientific, Rockford, IL, USA). A sample of 10 frozen eggs was dissolved in 4 mL of NaOH (0.1 M) in a 60°C water bath. Due to the high protein content of the eggs, the protein subsamples were diluted (1:100) with NaOH (0.1 M) before the analysis. The protein content of the diluted subsamples was quantified using a Shimadzu UV Mini-2040 UV-VIS spectrophotometer at 595 nm. The average protein content of an individual egg (mg * wet weight^-1^) was calculated as the average of two samples standardized with the known protein concentration of bovine serum albumin. Standardization was done using a linear curve. Egg lipid to protein ratio was calculated as total egg lipid content divided by the sum of egg lipid and protein content.

### Statistical analyses

All statistical analyses were conducted in the *R* environment version 4.2.1 (R Core Team, 2022). The effect of diet and population on female maturation was assessed with a χ^2^ test. In the maturation rate analyses, females that were stripped were considered as mature and the remaining fish as immature. 59 mature and 741 immature females were included in the analysis on the effect of diet on maturation rate. Population information was missing for four of the mature and seven of the immature females. These females were thus removed from the analysis on the effect of population on maturation bringing the final number of mature and immature females included in the analysis to 55 and 734, respectively.

A total of 49 females were included in the final female and egg trait analyses after removing the females that were stripped at a sub-optimal time (*n* = 2) and females with missing egg trait data (*n* = 4) or population and family information (*n* = 4). To analyse the effect of female diet on female and egg traits, linear mixed effects models (LMM) were fitted using the *lme4* version 1.1-30 package (Bates *et al*., 2015). All models were first fitted with second- and third-degree interactions between the explanatory variables but as these were found to not be significant (α = 0.05), the interactions were removed from subsequent analyses. All models included tank and parental IDs (sires and dams) as random intercepts to account for non-independence between the samples.

The mature female traits that were analysed included body weight prior to egg stripping, body length, condition factor and fecundity (number of eggs stripped). Feed treatment was included as an explanatory variable in all of the models. Female body size can influence fecundity (Barneche *et al*., 2018) and therefore, female body weight prior to egg stripping was also added as a fixed effect in the fecundity model. Body weight was chosen as an explanatory variable over body length or condition because weight tends to explain more of the variation in fecundity compared to length (Koops *et al*., 2004). Moreover, female condition should be measured months prior to spawning (Koops *et al*., 2004) as gonad development takes place during a period of fasting and accumulated energy stores are utilized throughout the reproductive season (Jonsson *et al*., 1991).

The egg traits that were analysed included egg size and egg lipid and protein content. The initial egg trait models included feed treatment and female body size at the time of stripping as explanatory variables. As the different egg size measurements (egg diameter and egg dry and wet weight) are correlated with one another (Table S3), we used principal components analysis (PCA) to detect patterns within the egg size variables. The majority of the variation (85.5%) was explained by the first principal component (PC1) where the variables were all highly correlated with one another. The second and third principal components explained only little of the variation (12.8% and 1.7%, respectively). Moreover, PC1 was the only principal component with an eigenvalue >1. Thus, PC1 was used as a proxy for egg size in the subsequent analyses. In addition to feed treatment and female body size, egg size (PC1) was included as an explanatory variable in the egg lipid and protein content models.

As female body size may affect egg traits (e.g., Barneche *et al*., 2018; Heinimaa & Heinimaa, 2004), we fitted three separate models for each egg trait analysed, and of each of those, three different female body size measurements (i.e., body weight, body length, or condition) used as a fixed effect. These three different models for each of the three egg traits were fitted using maximum-likelihood estimates and compared using Akaike information criterion corrected for small sample sizes (AICc) (Burnham & Anderson, 2002). The differences in AICc values between the models were small (ΔAICc < 2) indicating similar predictive power for all the models. We present female body weight as an explanatory variable in the final egg trait models as the models with body weight were found to consistently have the lowest or second lowest AICc score of the compared models (Table S4). The final models were re-fitted using restricted maximum-likelihood (REML) estimates.

Means and one standard error of the mean (± SE) are reported for the different female and egg traits. Egg lipid content was natural log (*ln*) transformed and used in the final model to improve normality of the residuals. Model fits were assessed by inspecting residual plots (i.e., normal Q-Q plots, standardized vs fitted residuals, and histograms) using the *performance* version 0.9.2 (Lüdecke *et al*., 2021) and *DHARMa* version 0.4.5 (Hartig, 2022) packages. Models did not show any clear violations of assumptions. 95% confidence intervals (95% CI) were calculated for all model estimates with parametric bootstrapping using 5 000 iterations. For the egg protein model, parametric bootstrapping failed to converge due to the zero variance associated with the model random effects. Therefore, additional confidence intervals were calculated using the same bootstrapping method for a simplified linear egg protein model without random effects. The 95% CI values from the linear model were nearly identical with the confidence intervals obtained for the original mixed effect model containing the tank and parental ID random effects. For the egg protein model, the reported confidence intervals are ones obtained from the simplified linear model. Data management and visualisation was done using the *tidyverse* version 1.3.2 package (Wickham *et al*., 2019).

### Ethical statement

The experiments were approved by the Project Authorisation Board (ELLA) on behalf of the Regional Administrative Agency for Southern Finland (ESAVI) under experimental license ESAVI/2778/2018.

## Results

### Maturity

Of the 800 female Atlantic salmon in the warm-water treatment, 59 matured during the 2020–2021 spawning season (7.4%, Table S5). None of the 1 059 females in the cold-water treatment matured and were therefore not included in further female or egg trait analyses.

We found marked differences in maturation rates of the two populations included in the study (*χ*^2^ = 29.82, df = 1, *p* = < 0.001; Fig. 1). Of the mature females, 44, originating from twenty different families, were from the Neva population and 11 (from three families) from the Oulu population (Table S5). The maturation rates of the Neva females were similar, 13.0% and 12.4%, in the control and low-fat feed treatments, respectively (Fig. 1). Oulu females had lower maturation rates than Neva females, but these rates were also relatively similar (2.3% and 2.6%) in the two feed treatments (Fig. 1). Due to the low number of Oulu females that matured compared to Neva, statistical analyses of female and egg traits with a reduced data set including only the Neva females were conducted. Removal of the Oulu females did not change the overall inference of the results (data not shown). Therefore, the two populations were pooled together for subsequent statistical analyses. Based on the pooled data, diet did not influence maturation rate and there were similar proportions of mature females in both feed treatments with 28 mature out of 383 and 31 mature out of 417 individuals in control and low-fat treatments, respectively (*χ*^2^ = 4.20e^-29^, df = 1, *p* = 1.00; Fig. 1).

**Fig. 1.**
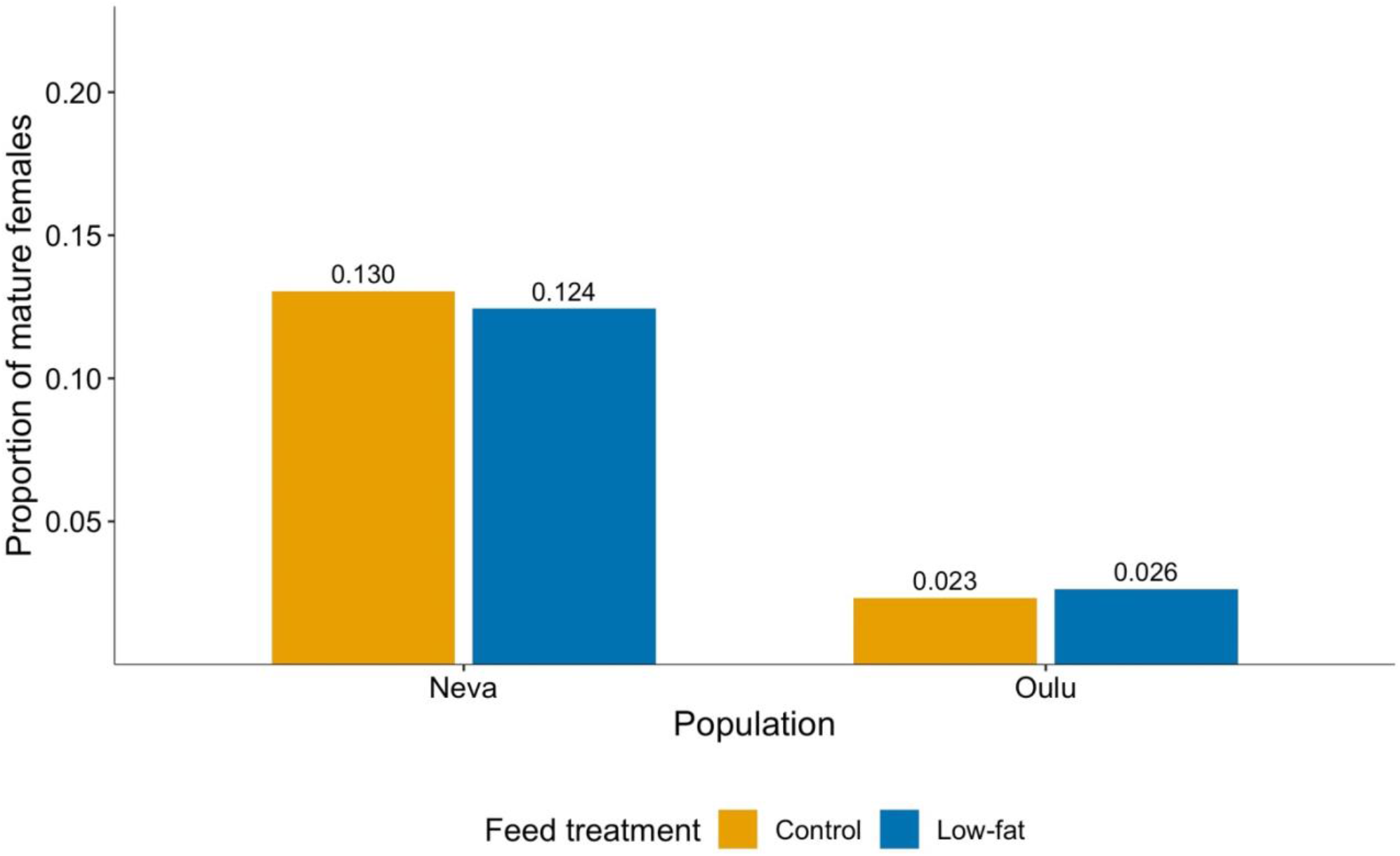
The proportion of mature female Atlantic salmon of the Neva (*n* = 44) and Oulu (*n* = 11) populations in the control (orange) and low-fat (blue) feed treatments. The value above the bar indicates the exact proportion of mature females found within each population and feed treatment.

### Female traits

At the time of the study, females in the cold-water treatment were, on average, smaller with a mean body weight of 0.41 ± 0.01 kg (mean ± SE) compared to the females in the warm-water treatment that had a mean body weight of 1.23 ± 0.02 kg (Fig. 2). The females in the warm-water treatment showed a slightly wider range of body size compared to the females in the cold-water treatment (Fig. 2). The body weight of the mature females ranged from 0.77 kg to 2.20 kg (Figs. 2 and S3). The mature females were slightly heavier with a mean weight of 1.39 ± 0.04 kg compared to the average weight of all the females in the warm-water treatment. The weight of the mature females was similar between the control and low-fat feed treatments (Tables 1 and 2). Similarly, the length and condition factor of the mature females was not influenced by the energy content of the diet (Table 2). Female fecundity varied little between the two feed treatments ranging from 1 321 to 6 885 eggs per female (Fig. S1) with mean fecundity in both feeding treatments being above 3 000 (Table 1). There was a significant, positive correlation between female body weight and fecundity (Table 2, Fig. S3), with every additional 100 g in female weight resulting in 256 ± 18 and 240 ± 12 additional eggs in the control and low-fat feed treatments, respectively. Female diet, however, did not influence the number of eggs mature females produced (Table 2).

**Table 1.** Female and egg traits of mature Atlantic salmon females in each feeding treatment. All values are presented as mean ± SE and egg traits as mg * wet weight^-1^. Relative fecundity indicates the number of eggs a female Atlantic salmon produces per 100 g of body wet mass. The relative amount of different egg components is given as % of egg wet weight. Other represents the residual weight after subtracting water, lipid, and protein from the egg wet weight. The table includes female and egg trait information on mature females that were included in the final statistical analyses.

|  | Feed treatment |  |
| --- | --- | --- |
|  | Control | Low-fat |
|  | *n* = 22 | *n* = 27 |
| **Female traits** |  |  |
| Weight (kg) | 1.5 ± 0.1 | 1.4 ± 0.1 |
| Length (cm) | 47.1 ± 0.7 | 45.7 ± 0.6 |
| Condition factor | 1.4 ± 0.0 | 1.4 ± 0.0 |
| Fecundity (egg *n*) | 3740 ± 257 | 3317 ± 208 |
| Relative fecundity (egg *n* *100 g<sup>-1</sup>) | 256 ± 18 | 240 ± 12 |
| **Egg traits** |  |  |
| Diameter (mm) | 4.5 ± 0.1 | 4.4 ± 0.0 |
| Wet weight (mg) | 46.4 ± 1.4 | 44.7 ± 1.0 |
| Dry weight (mg) | 16.3 ± 0.5 | 14.2 ± 0.4 |
| Lipid (mg) | 2.4 ± 0.1 | 1.6 ± 0.1 |
| Protein (mg) | 1.6 ± 0.1 | 1.6 ± 0.1 |
| Water (mg) | 30.2 ± 1.1 | 30.5 ± 0.8 |
| Other (mg) | 12.3 ± 0.4 | 11.0 ± 0.4 |
| %Lipid | 5.2 ± 0.2 | 3.6 ± 0.1 |
| %Protein | 3.5 ± 0.1 | 3.7 ± 0.2 |
| %Water | 64.8 ± 0.6 | 68.1 ± 0.8 |
| %Other | 26.5 ± 0.5 | 24.7 ± 0.7 |

**Table 2.**
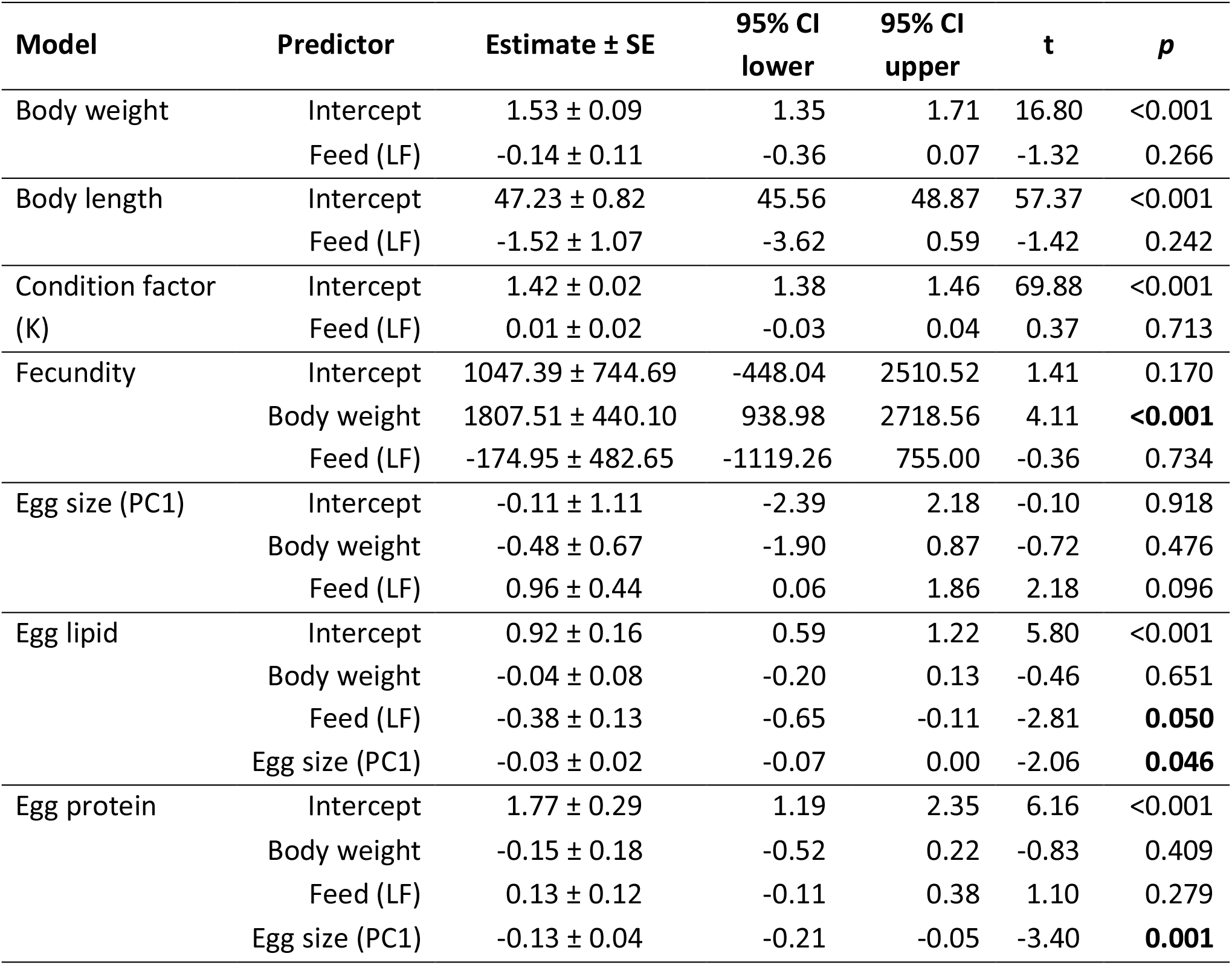
Results of the linear mixed models for female and egg traits of mature Atlantic salmon females. Estimates are presented as treatment contrasts with control feed treatment as the intercept. The estimate 95% confidence intervals were calculated using parametric bootstrapping with 5 000 iterations. Bolded *p*-values indicate statistical significance at a 0.05 level. LF = low-fat feed treatment.

**Fig. 2.**
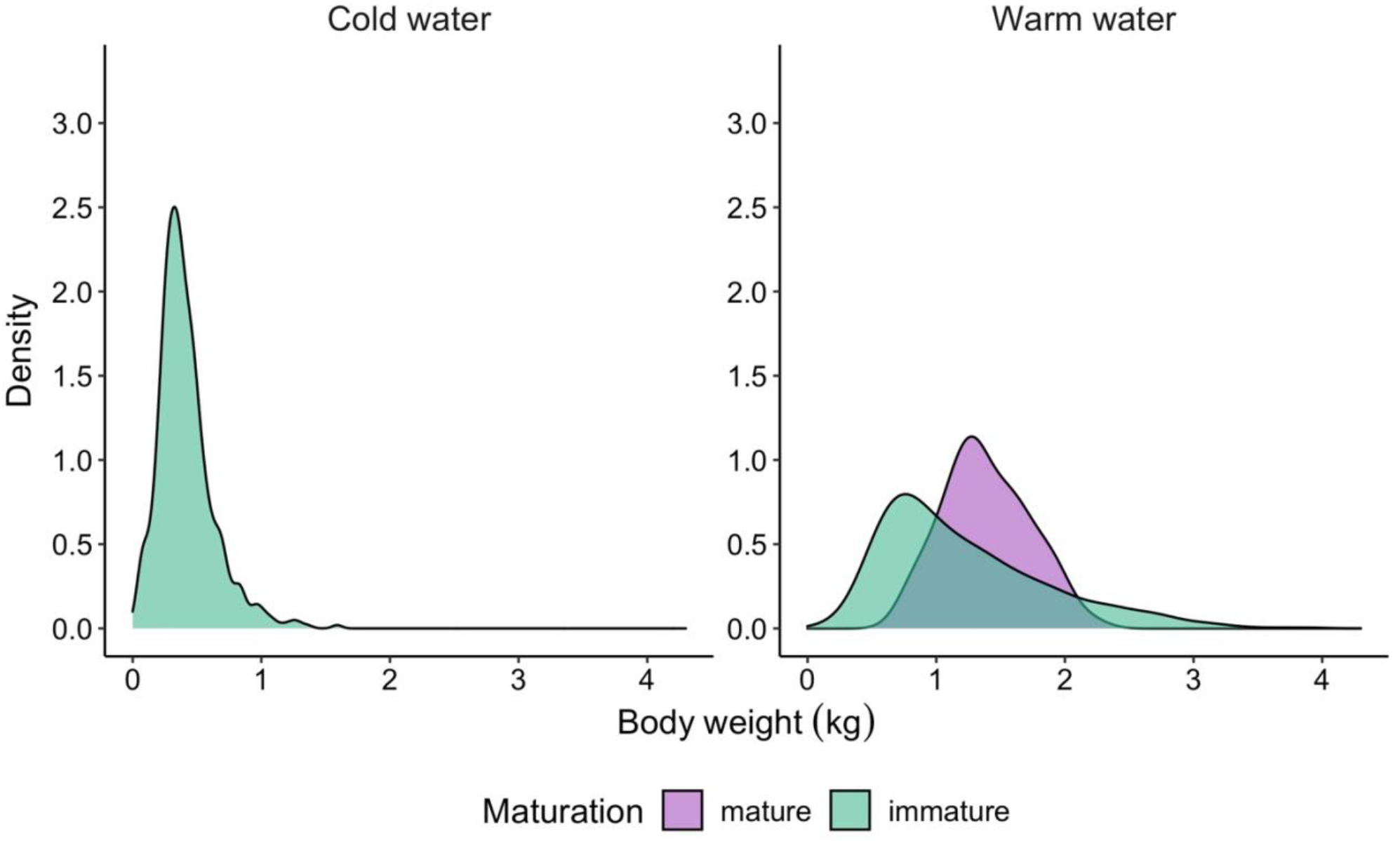
Body weight distribution of mature (purple) and immature (green) female Atlantic salmon during the 2020–2021 spawning season in the cold-water (left panel) and warm-water (right panel) treatments. There were a total of 1059 females in the cold-water treatment none of which matured. In the warm-water treatment, there were 800 females of which 59 matured during the spawning season.

### Egg traits

For females in the control feed treatment, an average egg was comprised of 64.8 ± 0.6% water, 5.2 ± 0.2% lipid, and 3.5 ± 0.1% protein (Table 1) compared to 68.1 ± 0.8% water, 3.6 ± 0.1% lipid, and 3.7 ± 0.2% protein in the low-fat feed treatment (Table 1). Egg size (PC1) was not significantly affected by either diet or body weight (Table 2, Fig. 3). Likewise, female body weight had no marked effect on egg lipid or protein content (Table 2). The amount of lipid and protein in the eggs was, however, influenced by egg size with smaller eggs containing less lipids and proteins (Table 2). The differences between the energy content of the female diet were reflected in the egg molecular composition. The total mean egg lipid content of the female salmon receiving the higher energy control diet was significantly greater at 2.4 ± 0.1 mg * wet weight^-1^ compared to the females on the low-fat diet that had a mean egg lipid content of 1.6 ± 0.1 mg * wet weight^-1^ (Tables 1 and 2, Fig. 3). On the other hand, feed treatment had no significant effect on protein content of the eggs (Table 2, Fig. 3). On average, the lipid to protein ratio of the eggs was greater for the females fed on the higher energy content control diet (Fig. 3).

**Fig. 3.**
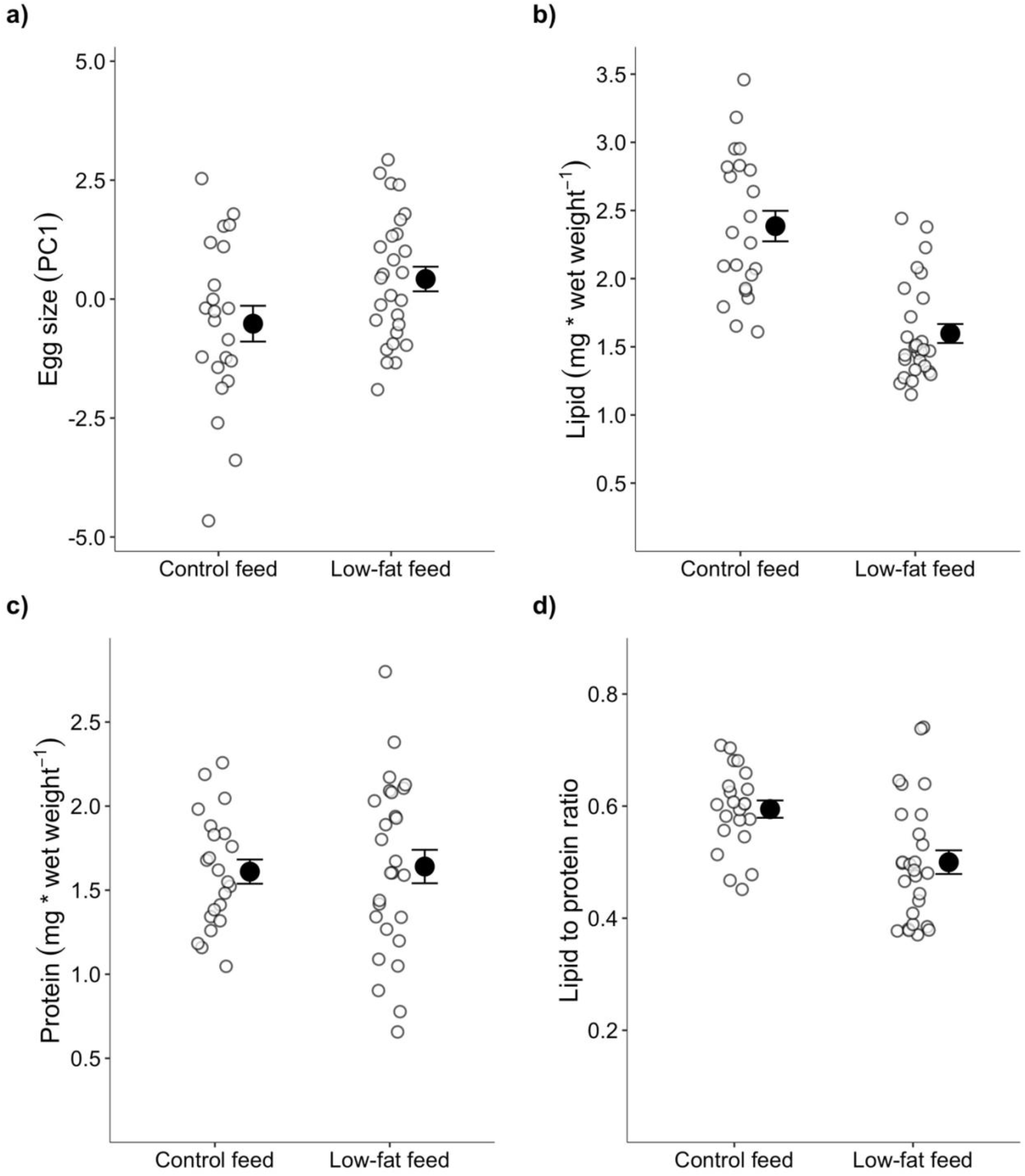
Egg a) size, b) lipid content, c) protein content, and d) lipid to protein ratio in the control and low-fat feed treatments. Smaller open circles represent individual measurements and larger filled circles with error bars represent mean ± SE.

## Discussion

In this study, we investigated how the environment experienced by maturing female Atlantic salmon influenced age of maturation, fecundity, and egg traits. We demonstrate that temperature affected female maturation rate at three years of age, whereas diet did not. However, diet affected egg traits. Specifically, a higher energy maternal diet resulted in greater egg lipid content compared to a 22% lower energy diet. These results suggest that female salmon with a lower energy diet may have reduced reproductive fitness if the diet has an adverse effect on the development, growth, and subsequent survival of the offspring via maternal egg provisioning.

This study was part of a larger experiment investigating the effect of temperature on sexual maturation in Atlantic salmon. A 2°C lower temperature was sufficient to fully delay maturation in female Atlantic salmon for at least one reproductive season. The females in the cold-water treatment were on average smaller and exhibited less variation in body weight compared to the females in the warm-water treatment. The absence of mature females in the cold-water treatment is likely due to reduced growth rate and lower rate of lipid accumulation and consequently lower somatic lipid stores at cooler temperatures (Jonsson & Jonsson, 2011; Thorpe, 1994). Mature females had likely acquired sufficient somatic energy reserves allowing for maturation whereas the immature females had not. Because no females matured in the cold-water treatment, we could not assess the effect temperature may have on egg traits. The Atlantic salmon in our study represented two different genetically distinct Finnish populations (Säisä *et al*., 2005). The two populations exhibited different rates of maturation with the females of the Neva population showing higher rates of maturation compared to the Oulu females. These population differences could, in part, be due to lower growth rate and smaller body size of the Oulu fish as has been observed for the males in this study population (Åsheim *et al*., 2022). Although the body size of the Oulu fish overall was smaller compared to the Neva population, we did not find significant differences between the populations in the size of the females that matured. The lack of body size differences in maturation may indicate that the two populations share a similar threshold for body size and accumulated energy store necessary for maturation in females (Thorpe *et al*., 1998).

Contrary to our expectations, the reduction in energy content of the diet initiated about 1.5 years before maturation assessments was not associated with female maturation rate in our study. These results are somewhat surprising as previous studies that have modified the energy content of the diet have shown clear associations between diet and maturation in salmonids for both males and females (Jonsson *et al*., 2013, 2012; Shearer & Swanson, 2000). The absence of a dietary effect on maturation is potentially due to the specific energy restriction strategy employed in our study. Rather than reducing food amount, we provided feed of lower energy content *ad libitum* in order to limit the potential for behavioural differences contributing to differences in feed intake. This may have resulted in the fish acquiring adequate lipid stores needed for maturation regardless of the energy content of their diet. Alternatively, the reduction in feed energy content in this experiment might not have been severe enough for any dietary effects on maturation to be detected.

It is well established that female body size and fecundity are highly correlated in fishes (Bagenal, 1969; de Eyto *et al*., 2015; Heinimaa & Heinimaa, 2004; Millner *et al*., 1991; Reid & Chaput, 2012; Smalås *et al*., 2017; Thorpe *et al*., 1984). A similar pattern was found in this study, where fecundity increased with female body size with an increase of approximately 7.4% in egg number for every 100 grams of added body weight. The females in our study produced more eggs per unit of body weight compared to some previous studies (e.g., de Eyto *et al*., 2015; Heinimaa & Heinimaa, 2004; Reid & Chaput, 2012). This difference is likely due to the mature females in our study being smaller (mostly under 2 kg) compared to the aforementioned studies (body weight range from ∼2 kg to up to 16 kg) as smaller Atlantic salmon females often have higher fecundity per unit of body size (Fleming, 1996).

Feed energy content did not affect female fecundity in our experiment. The effects of diet on salmonid fecundity vary as there are multiple opportunities to alter the fish’s diet ranging from reducing the feed amount to making various nutritional changes, and such studies are showing varying results on fecundity. Bagenal (1969) and Scott (1962) found that decreasing the ration size led to a decrease in fecundity in brown trout (*Salmo trutta*) and rainbow trout (*Oncorhynchus mykiss*) even after taking the female body size into account. The importance of ration size on fecundity has also been demonstrated for many non-salmonid species of fish (reviewed in Fernández-Palacios *et al*., 2011). Diet studies on Atlantic salmon and rainbow trout that have altered the micronutrient or lipid composition of the female diet, on the other hand, have shown an effect on fecundity in some (Eskelinen, 1989; Washburn *et al*., 1990) but not all cases (Rennie *et al*., 2005; Sandnes *et al*., 1984; Yıldız *et al*., 2020). The effect diet may have on female fecundity appears to vary not only between species but also between the diets’ nutritional composition (reviewed in Fernández-Palacios *et al*., 2011). The results from our study and the varied results from previous dietary nutritional content and ration size studies indicate that feed intake may have a greater influence on female fecundity than the feed’s specific nutritional composition.

In this experiment, we measured a variety of egg traits to understand how diet may influence maternal provisioning of offspring via egg size and nutritional composition. Diet did not affect egg size in our study. This is in line with previous studies that have also failed to observe an effect of the female’s diet’s nutritional composition on egg size in Atlantic salmon and rainbow trout (Eskelinen, 1989; Rennie *et al*., 2005; Sandnes *et al*., 1984; Yıldız *et al*., 2020). In contrast, we expected to see an effect of dietary energy content on egg molecular composition as nutrients are allocated to the eggs from the maternal somatic resources during gonad development (Brooks *et al*., 1997; Wootton & Smith, 2015). In our study, reduction in dietary energy content was reflected in the lipid content of the eggs where the eggs from females fed the low-fat diet produced eggs with lower lipid content. Egg protein content, on the other hand, was not significantly affected by diet (though estimated as somewhat higher under the low-fat diet) which led to a higher lipid-to-protein ratio in the eggs in the control feed treatment, on average. Both egg lipid and protein content were influenced by egg size with larger eggs holding a greater amount of lipid and protein. Decrease in dietary quality in terms of energy content may lead to a female salmon having lower somatic energy stores from which to allocate resources to their eggs without compromising their own survival (Jonsson & Jonsson, 2011; Klemetsen *et al*., 2003). Moreover, lipids and proteins present in eggs are a crucial energy source and a material source for biosynthesis of various molecules for the developing embryos and alevins (hatched juveniles with yolk still attached) (Berg *et al*., 2001; Brooks *et al*., 1997; Finn & Fyhn, 2010; Kamler, 2008; Thorstad *et al*., 2010). For example, alevins with greater energy stores may have lower starvation-induced mortality and an initial growth advantage over alevins whose mothers are unable to provide them with higher energy resources (Berg *et al*., 2001; Einum, 2003; Ojanguren *et al*., 1996; Thorn *et al*., 2019). Thus, our results suggest that female salmon experiencing a lower energy diet may experience a potential reproductive fitness disadvantage via lower egg energy provisioning of the offspring in the form of lipid content.

Female body size or condition did not affect egg composition in our study. Previously, larger Atlantic salmon females have been shown to produce eggs of higher energy content in a wild population (Heinimaa & Heinimaa, 2004). In the study by Heinimaa and Heinimaa (2004), the mature females were from a wild population and represented a wide range of body sizes (∼5.0 – 16.0 kg). The size range of the mature females in our study, on the other hand, was much narrower (∼0.80 – 2.20 kg) and the salmon represented F2 generation hatchery individuals. In addition, the previous study compared across age classes whereas we compared within a common age class. Any of these differences could potentially explain why we were unable to find an effect of female body size on egg composition.

Possible genetic effects on female reproductive traits influencing the results of our study should not be ruled out. For example, the *vgll3* locus has been identified as affecting maturation age in Atlantic salmon (Ayllon *et al*., 2015; Barson *et al*., 2015). Similarly, a recent study demonstrated that the *six6* genomic region, which is also associated with maturation age in males (Sinclair-Waters *et al*., 2020), associates with stomach fullness in wild Atlantic salmon (Aykanat *et al*., 2020). Thus, genetic effects may contribute to female reproductive traits as the traits are dependent on female resource acquisition, the consequent somatic energy reserves, and the utilization of these energy reserves.

Understanding how environmental variables shape reproductive traits is a fundamental role of evolutionary ecology. Our results provide evidence for the link between two key environmental factors females experience and the potential quality of their eggs. Additionally, the differences in maturation rates between the two temperatures observed in this study may have implications for Atlantic salmon population dynamics if warmer water temperatures experienced due to climate change lead to earlier maturation ages and thus shorter generation times. In the future, multi-generation experiments combining measurements of egg nutritional content, fertilization, hatching success, and juvenile survival rates while taking genetic and environmental factors into consideration would further our knowledge on salmon reproduction and the role parental effects play in offspring fitness.

## Acknowledgements

We wish to thank the many people who’ve participated in this project over the years. We thank Jacqueline Moustakas-Verho, Ksenia Zueva, Marion Sinclair-Waters, Nico Lorenzen, Victoria Pritchard, Yann Czorlich, Spiros Papakostas, and Suvi Ikonen for their help with gamete collection and performing the crosses that created the salmon in this study. Thank you to Natural Resources Institute Finland (LUKE) for access to the broodstocks and staff of the Taivalkoski and Laukaa hatcheries (LUKE) for their help with gamete collection. Nikolai Piavchenko and Noora Parre are thanked for animal husbandry at the Viikki campus. We thank Lammi Biological Station for hosting our common garden facilities and especially John Loehr for assistance with local logistics as well as Susanna Airaksinen and Thomas Ginström and others at Raisioaqua Ltd for producing the feed used in the study. We are grateful to Suvi Ikonen, Andrés Salgado, Anna Toikkanen, Dorian Jagusch, Fin Morrison, Ike van Gestel, Jeferson Delgado Florez, Maël Le Gouellec, Markus Lauha, Mikko Immonen, Paul Bangura, Teemu Mäkinen, and Tiemen Jansen for their assistance with fish husbandry at Lammi Biological Station. J. Delgado Florez, M. Le Gouellec, M. Lauha, M. Immonen, P. Bangura, and S. Ikonen are thanked for their assistance with egg sampling and Iikki Donner for technical assistance with lipid extractions. We thank Annukka Ruokolainen, Seija Tillanen, Shadi Jansouz, I. Donner, Morgane Frapin, and Tutku Aykanat for sex determination and population and parentage assignment of the fish.

## Funding

This project received funding from the European Research Council (ERC) under the European Union’s Horizon 2020 research and innovation programme (grant agreement No 742312) and from the Academy of Finland grants 307593, 302873 and 284941 to C.R.P. and from Societas pro Fauna et Flora Fennica to K.S.M.

## Author contributions

K.S.M., C.R.P., & K.B.M. conceived the study. P.V.D. & C.R.P. designed the broodstock crosses. J.E. provided access to salmon broodstocks. P.L., E.R.Å., A.H.H., & K.S.M. coordinated fish husbandry. C.R.P. coordinated the molecular data generation. K.S.M., E.R.Å., C.R.P., & K.B.M. coordinated sampling. K.S.M. analysed the data. K.S.M. drafted the manuscript with input from all authors.

## Conflict of interest

The authors declare no conflict of interest.

## Data and materials availability

All data and code will be provided via Zenodo upon acceptance of the article.

## Supplementary material

Tables S1 – S5 and Figures S1 – S3.

## Notes

### Competing Interest Statement

The authors have declared no competing interest.

## References

Åsheim, E. R., Debes, P. V., House, A., Niemelä, P. T., Siren, J. P., Erkinaro, J., & Primmer, C. R. (2022). Strong effects of temperature, population and age-at-maturity genotype on maturation probability for Atlantic salmon in a common garden setting. bioRxiv 2022, doi:10.1101/2022.07.22.501167.

Aykanat, T., Rasmussen, M., Ozerov, M., Niemelä, E., Paulin, L., Vähä, J.-P., … Primmer, C. R. (2020). Life-history genomic regions explain differences in Atlantic salmon marine diet specialization. Journal of Animal Ecology, 89, 2677–2691.

Ayllon, F., Kjærner-Semb, E., Furmanek, T., Wennevik, V., Solberg, M. F., Dahle, G., … Wargelius, A. (2015). The vgll3 Locus Controls Age at Maturity in Wild and Domesticated Atlantic Salmon (Salmo salar L.) Males. PLoS Genetics, 11, 1–15.

Bagenal, T. B. (1969). The Relationship Between Food Supply and Fecundity in Brown Trout Salmo trutta L. Journal of Fish Biology, 1, 167–182.

Barneche, D. R., Robertson, D. R., White, C. R., & Marshall, D. J. (2018). Fish reproductive-energy output increases disproportionately with body size. Science, 360, 642–645.

Barson, N. J., Aykanat, T., Hindar, K., Baranski, M., Bolstad, G. H., Fiske, P., … Primmer, C. R. (2015). Sex-dependent dominance at a single locus maintains variation in age at maturity in salmon. Nature, 528, 405–408.

Bates, D., Mächler, M., Bolker, B. M., & Walker, S. C. (2015). Fitting linear mixed-effects models using lme4. Journal of Statistical Software, 67.

Berg, O. K., Hendry, P., Svendsen, B., Bech, C., Arnekleiv, J. V., & Lohrmann, A. (2001). Maternal provisioning of offspring and the use of those resources during ontogeny: Variation within and between Atlantic Salmon families. Functional Ecology, 15, 13–23.

Bogevik, A. S., Natário, S., Karlsen, O., Thorsen, A., Hamre, K., Rosenlund, G., & Norberg, B. (2012). The effect of dietary lipid content and stress on egg quality in farmed Atlantic cod Gadus morhua. Journal of Fish Biology, 81, 1391–1405.

Bradford, M. M. (1976). A Rapid and Sensitive Method for the Quantitation Microgram Quantities of Protein Utilizing the Principle of Protein-Dye Binding. Analytical Biochemistry, 72, 248–254.

Brooks, S., Tyler, C. R., & Sumpter, J. P. (1997). Egg quality in fish: What makes a good egg? Reviews in Fish Biology and Fisheries, 7, 387–416.

Burnham, K. P., & Anderson, D. R. (2002). Model selection and multimodel inference: a practical information-theoretic approach, 2nd ed. Springer New York.

Debes, P. V., Piavchenko, N., Erkinaro, J., & Primmer, C. R. (2020). Genetic growth potential, rather than phenotypic size, predicts migration phenotype in Atlantic salmon: Determinants of migration phenotypes. Proceedings of the Royal Society B: Biological Sciences, 287.

Duston, J., & Saunders, R. L. (1999). Effect of winter food deprivation on growth and sexual maturity of Atlantic salmon (Salmo salar) in seawater. Canadian Journal of Fisheries and Aquatic Sciences, 56, 201–207.

Einum, S. (2003). Atlantic salmon growth in strongly food-limited environments: Effects of egg size and paternal phenotype? Environmental Biology of Fishes, 67, 263–268.

Einum, S., & Fleming, I. A. (2000). Selection against late emergence and small offspring in Atlantic salmon (Salmo salar). Evolution, 54, 628–639.

Eskelinen, P. (1989). Effects of different diets on egg production and egg quality of Atlantic salmon (Salmo salar L.). Aquaculture, 79, 275–281.

de Eyto, E., White, J., Boylan, P., Clarke, B., Cotter, D., Doherty, D., … O’Higgins, K. (2015). The fecundity of wild Irish Atlantic salmon Salmo salar L. and its application for stock assessment purposes. Fisheries Research, 164, 159–169.

Fernández-Palacios, H., Nordberg, B., Izquierdo, M., & Hamre, K. (2011). Effects of broodstock diet on eggs and larvae. In G. J. Holt (Ed.), Larval Fish Nutrition (pp. 153–181). John Wiley & Sons, Ltd.

Finn, R. N., & Fyhn, H. J. (2010). Requirement for amino acids in ontogeny of fish. Aquaculture Research, 41, 684–716.

Fjelldal, P. G., Hansen, T., & Huang, T. (2011). Continuous light and elevated temperature can trigger maturation both during and immediately after smoltification in male Atlantic salmon (Salmo salar). Aquaculture, 321, 93–100.

Fleming, I. A. (1996). Reproductive strategies of Atlantic salmon: Ecology and evolution. Reviews in Fish Biology and Fisheries, 6, 379–416.

Gagliano, M., & McCormick, M. I. (2007). Maternal Condition Influences Phenotypic Selection on Offspring. Journal of Animal Ecology, 76, 174–182.

Green, B. S. (2008). Chapter 1 - Maternal Effects in Fish Populations. In D. W. Sims (Ed.), Advances in Marine Biology (pp. 1–105). Academic Press.

Gregersen, F., Vøllestad, L. A., Ø stbye, K., Aass, P., & Hegge, O. (2011). Temperature and food-level effects on reproductive investment and egg mass in vendace, Coregonus albula. Fisheries Management and Ecology, 18, 263–269.

Hartig, F. (2022). DHARMa: Residual Diagnostics for Hierarchical (Multi-Level / Mixed) Regression Models.

Heincke, F. (1908). Bericht über die Untersuchungen der Biologischen Anstalt auf Helgoland zur Naturgeschichte der Nutzfische. IV/V. Bericht über die Beteiligung Deutschlands an der Internationalen Meeresforschung in den Jahren 1905/6–1906/7. Berlin. pp. 67–156.

Heinimaa, S., & Heinimaa, P. (2004). Effect of the female size on egg quality and fecundity of the wild Atlantic salmon in the sub-arctic River Teno. Boreal Environment Research, 9, 55–62.

Imsland, A. K., Handeland, S. O., & Stefansson, S. O. (2014). Photoperiod and temperature effects on growth and maturation of pre- and post-smolt Atlantic salmon. Aquaculture International, 22, 1331–1345.

Izquierdo, M. S., Fernández-Palacios, H., & Tacon, A. G. J. (2001). Effect of broodstock nutrition on reproductive performance of fish. Aquaculture, 197, 25–42.

Jonsson, B., Jonsson, N., & Finstad, A. G. (2013). Effects of temperature and food quality on age and size at maturity in ectotherms: an experimental test with Atlantic salmon. Journal of Animal Ecology, 82, 201–210.

Jonsson, B., & Jonsson, N. (2011). Ecology of Atlantic Salmon and Brown Trout - Habitat as a Template for Life Histories. Springer.

Jonsson, B., Finstad, A. G., & Jonsson, N. (2012). Winter temperature and food quality affect age at maturity: An experimental test with Atlantic salmon (Salmo salar). Canadian Journal of Fisheries and Aquatic Sciences, 69, 1817–1826.

Jonsson, N., Jonsson, B., & Hansen, L. P. (1991). Energetic cost of spawning in male and female Atlantic salmon (Salmo salar L.). Journal of Fish Biology, 39, 739–744.

Jonsson, N., & Jonsson, B. (2003). Energy allocation among developmental stages, age groups, and types of Atlantic salmon (Salmo salar) spawners. Canadian Journal of Fisheries and Aquatic Sciences, 60, 506–516.

Kadri, S., Mitchell, D. F., Metcalfe, N. B., Huntingford, F. A., & Thorpe, J. E. (1996). Differential patterns of feeding and resource accumulation in maturing and immature Atlantic salmon, Salmo salar. Aquaculture, 142, 245–257.

Kamler, E. (1992). Early Life History of Fish: An energetics approach, 1st ed. Chapman & Hall, London.

Kamler, E. (2008). Resource allocation in yolk-feeding fish. Reviews in Fish Biology and Fisheries, 18, 143–200.

Kjesbu, O. S., Witthames, P. R., Solemdal, P., & Greer Walker, M. (1998). Temporal variations in the fecundity of Arcto-Norwegian cod (Gadus morhua) in response to natural changes in food and temperature. Journal of Sea Research, 40, 303–321.

Kjørsvik, E., Mangor-Jensen, A., & Holmefjord, I. (1990). Egg quality in fishes. Advances in Marine Biology, 26, 71–113.

Klemetsen, A., Amundsen, P. A., Dempson, J. B., Jonsson, B., Jonsson, N., O’Connell, M. F., & Mortensen, E. (2003). Atlantic salmon Salmo salar L., brown trout Salmo trutta L. and Arctic charr Salvelinus alpinus (L.): A review of aspects of their life histories. Ecology of Freshwater Fish, 12, 1–59.

Koops, M. A., Hutchings, J. A., & McIntyre, T. M. (2004). Testing hypotheses about fecundity, body size and maternal condition in fishes. Fish and Fisheries, 5, 120–130.

Lüdecke, D., Ben-Shachar, M., Patil, I., Waggoner, P., & Makowski, D. (2021). performance: An R Package for Assessment, Comparison and Testing of Statistical Models. Journal of Open Source Software, 6, 3139.

Mazorra, C., Bruce, M., Bell, J. G., Davie, A., Alorend, E., Jordan, N., … Bromage, N. (2003). Dietary lipid enhancement of broodstock reproductive performance and egg and larval quality in Atlantic halibut (Hippoglossus hippoglossus). Aquaculture, 227, 21–33.

Millner, R., Whiting, C., Walker, M. G., & Witthames, P. (1991). Growth increment, condition and fecundity in sole (Solea solea (L.)) from the north sea and eastern english channel. Netherlands Journal of Sea Research, 27, 433–439.

Mobley, K. B., Aykanat, T., Czorlich, Y., House, A., Kurko, J., Miettinen, A., … Primmer, C. R. (2021). Maturation in Atlantic salmon (Salmo salar, Salmonidae): a review of ecological, genetic, and molecular processes. Rev Fish Biol Fisheries, 31, 523–571.

Ojanguren, A. F., Reyes-Gavilán, F. G., & Braña, F. (1996). Effects of egg size on offspring development and fitness in brown trout, Salmo gairdneri L. Aquaculture, 147, 9–20.

R Core Team. (2022). R: A language and environment for statistical computing. R Foundation for Statistical Computing. Vienna, Austria.

Reid, J. E., & Chaput, G. (2012). Spawning history influence on fecundity, egg size, and egg survival of Atlantic salmon (Salmo salar) from the Miramichi River, New Brunswick, Canada. ICES Journal of Marine Science ICES, 69, 1678–1685.

Rennie, S., Huntingford, F. A., Loeland, A.-L., & Rimbach, M. (2005). Long term partial replacement of dietary fish oil with rapeseed oil; Effects on egg quality of Atlantic salmon Salmo salar. Aquaculture, 248, 135–146.

Rowe, D. K., Thorpe, J. E., & Shanks, A. M. (1991). Role of Fat Stores in the Maturation of Male Atlantic Salmon (Salmo salar) Parr. Canadian Journal of Fisheries and Aquatic Sciences, 48, 405–413.

Rueden, C. T., Schindelin, J., Hiner, M. C., DeZonia, B. E., Walter, A. E., Arena, E. T., & Eliceiri, K. W. (2017). ImageJ2: ImageJ for the next generation of scientific image data. BMC Bioinformatics, 18, 529.

Säisä, M., Koljonen, M.-L., Gross, R., Nilsson, J., Tähtinen, J., Koskiniemi, J., & Vasemägi, A. (2005). Population genetic structure and postglacial colonization of Atlantic salmon (Salmo salar) in the Baltic Sea area based on microsatellite DNA variation. Canadian Journal of Fisheries and Aquatic Sciences, 62, 1887–1904.

Sandnes, K., Ulgenes, Y., Braekkan, O. R., & Utne, F. (1984). The effect of ascorbic acid supplementation in broodstock feed on reproduction of rainbow trout (Salmo gairdneri). Aquaculture, 43, 167–177.

Scott, D. P. (1962). Effect of Food Quantity on Fecundity of Rainbow Trout, Salmo gairdnerit. Journal of the Fisheries Research Board of Canada, 14, 715–731.

Shearer, K. D., & Swanson, P. (2000). The effect of whole body lipid on early sexual maturation of 1 + age male chinook salmon (Oncorhynchus tshawytscha). Aquaculture, 190, 343–367.

Shearer, K., Parkins, P., Gadberry, B., Beckman, B., & Swanson, P. (2006). Effects of growth rate/body size and a low lipid diet on the incidence of early sexual maturation in juvenile male spring Chinook salmon (Oncorhynchus tshawytscha). Aquaculture, 252, 545–556.

Shinn, S. E., & Proctor, A. (2013). Rapid lipid extraction from egg yolks. JAOCS, Journal of the American Oil Chemists’ Society, 90, 315–316.

Sinclair-Waters, M., Ødegård, J., Korsvoll, S. A., Moen, T., Lien, S., Primmer, C. R., & Barson, N. J. (2020). Beyond large-effect loci: Large-scale GWAS reveals a mixed large-effect and polygenic architecture for age at maturity of Atlantic salmon. Genetics Selection Evolution, 52, 1–11.

Sink, T. D., & Lochmann, R. T. (2008). Effects of dietary lipid source and concentration on channel catfish (Ictalurus punctatus) egg biochemical composition, egg and fry production, and egg and fry quality. Aquaculture, 283, 68–76.

Smalås, A., Amundsen, P. A., & Knudsen, R. (2017). The trade-off between fecundity and egg size in a polymorphic population of Arctic charr (Salvelinus alpinus (L.)) in Skogsfjordvatn, subarctic Norway. Ecology and Evolution, 7, 2018–2024.

Srivastava, R. K., & Brown, J. A. (1991). The biochemical characteristics and hatching performance of cultured and wild Atlantic salmon (Salmo salar) eggs. Canadian Journal of Zoology, 69, 2436–2441.

Taranger, G. L., Carrillo, M., Schulz, R. W., Fontaine, P., Zanuy, S., Felip, A., … Hansen, T. (2010). Control of puberty in farmed fish. General and Comparative Endocrinology, 165, 483–515.

Thorn, M. W., Dick, M. F., Oviedo, L., Guglielmo, C. G., & Morbey, Y. E. (2019). Transgenerational effects of egg nutrients on the early development of Chinook salmon (Oncorhynchus tshawytscha) across a thermal gradient. Canadian Journal of Fisheries and Aquatic Sciences, 76, 1253–1262.

Thorpe, J. E. (1994). Reproductive strategies in Atlantic salmon, Salmo salar L. Aquaculture Research, 25, 77–87.

Thorpe, J. E., Mangel, M., Metcalfe, N. B., & Huntingford, F. A. (1998). Modelling the proximate basis of salmonid life-history variation, with application to Atlantic salmon, Salmo salar L. Evolutionary Ecology, 12, 581–599.

Thorpe, J. E., Miles, M. S., & Keay, D. S. (1984). Developmental rate, fecundity and egg size in Atlantic salmon, Salmo salar L. Aquaculture, 43, 289–305.

Thorstad, E. B., Whoriskey, F., Rikardsen, A. H., & Aarestrup, K. (2010). Aquatic Nomads: The Life and Migrations of the Atlantic Salmon. In Ø. Aas, S. Einum, A. Klemetsen, & J. Skurdal (Eds.), Atlantic Salmon Ecology (pp. 1–32). Oxford, UK: Wiley-Blackwell.

Washburn, B. S., Frye, D. J., Hung, S. S. O., Doroshov, S. I., & Conte, F. S. (1990). Dietary effects on tissue composition, oogenesis and the reproductive performance of female rainbow trout (Oncorhynchus mykiss). Aquaculture, 90, 179–195.

Wickham, H., Averick, M., Bryan, J., Chang, W., McGowan, L. D., François, R., … Yutani, H. (2019). Welcome to the Tidyverse. Journal of Open Source Software, 4, 1686.

Wootton, R. J., & Smith, C. (2015). Reproductive Biology of Teleost Fishes. John Wiley & Sons, Ltd.

Yıldız, M., Ofori-Mensah, S., Arslan, M., Ekici, A., Yamaner, G., Baltacı, M. A., … Korkmaz, F. (2020). Effects of different dietary oils on egg quality and reproductive performance in rainbow trout Oncorhynchus mykiss. Animal Reproduction Science, 221, 106545.

Yoneda, M., & Wright, P. J. (2005). Effects of varying temperature and food availability on growth and reproduction in first-time spawning female Atlantic cod. Journal of Fish Biology, 67, 1225–1241.

