## Supplementary material for "Temperature and dietary energy content influence female maturation age and egg nutritional content in Atlantic salmon"

**This file includes:**

Fig. S1 – Fig. S3

Tables S1–S5

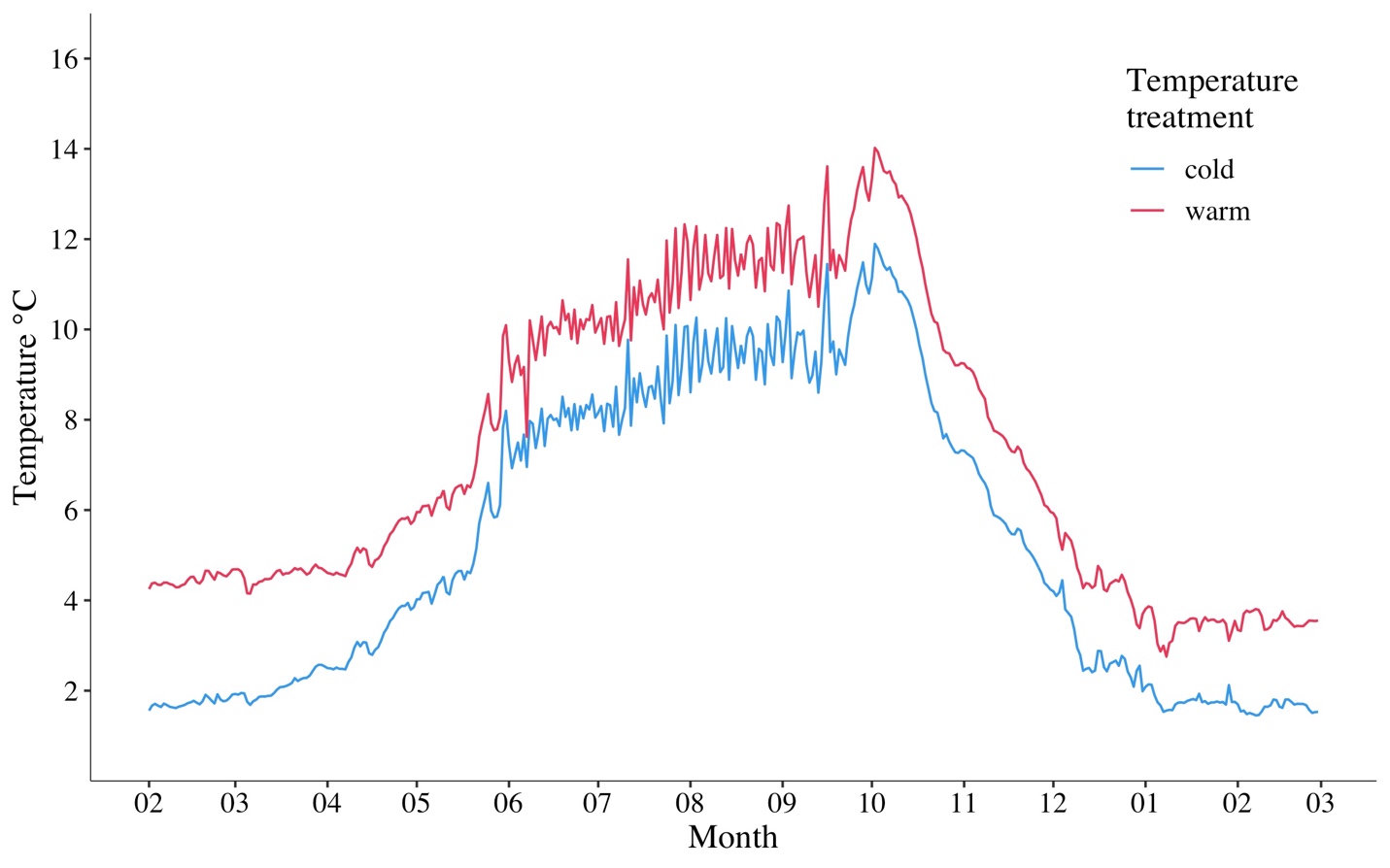

**Fig. S1**. Temperature variation in the experimental tanks at Lammi Biological Station between February 2020 and March 2021. Water temperature followed the natural temperature fluctuation of lake Pääjärvi from which water was pumped into the experimental tanks and a temperature difference of 2°C was created between the tanks with six tanks receiving warmed water and the other six cooled water.

**Fig. S2.** Timeline (not to scale) of the Atlantic salmon maturation experiment at the Lammi Biological Station (LBS). The timeline includes the major procedures relevant to this study on female maturation and egg traits during the spawning season of 2020/2021.

**Table S1**. Description of the three sampling periods of the study.

| **Sampling period** | **Date** | **Procedure** |
| --- | --- | --- |
| 1 | 2020-11-18 – 2020-12-02 | All females were assessed for signs of maturity and their maturity stage (I – IV) was recorded. Mature females were stripped. |
| 2 | 2021-01-04 – 2021-01-08 | All stage III females from sampling period 1 were checked and mature females were stripped. A part of the stage II females were checked for signs of maturity. |
| 3 | 2021-02-01 – 2021-02-17 | All females were checked for signs of maturity. Newly mature females were stripped and previously mature females were re-stripped if they were found to have new ripe eggs. |

**Table S2.** Description of maturation stages used to assess female maturity based on external characteristics.

| **Stage** | **Description** |
| --- | --- |
| I | No signs of maturity (II-IV) |
| II | Female has a rounded belly, especially towards the anterior end of the body |
| III | Female has a soft belly and/or extended cloaca |
| IV | Eggs are released when stripping |

**Table S3**. Pairwise Pearson’s correlation coefficients between the three different measures of egg size which were used in the principal components analysis.

|  | **Dry weight** | **Wet weight** |
| --- | --- | --- |
| **Egg diameter** | 0.721 | 0.947 |
| **Dry weight** |  | 0.670 |

**Table S4**. Egg trait LMM comparisons using AICc between fecundity and egg trait models that differ in female body size measurement used as a covariate.

| **Model** | **AICc** | Δ**AICc** |
| --- | --- | --- |
| Egg size ~ body length + feed + (1\|tank) + (1\|ID ma) + (1\|ID pa) | 188.97 |  |
| Egg size ~ body weight + feed + (1\|tank) + (1\|ID ma) + (1\|ID pa) | 189.05 | 0.08 |
| Egg size ~ K + feed + (1\|tank) + (1\|ID ma) + (1\|ID pa) | 189.32 | 0.27 |
| Egg lipid ~ body weight + feed + egg size + (1\|tank) + (1\|ID ma) + (1\|ID pa) | -9.08 |  |
| Egg lipid ~ body length + feed + egg size + (1\|tank) + (1\|ID ma) + (1\|ID pa) | -9.01 | 0.07 |
| Egg lipid ~ K + feed + egg size + (1\|tank) + (1\|ID ma) + (1\|ID pa) | -8.91 | 0.17 |
| Egg protein ~ K + feed + egg size + (1\|tank) + (1\|ID ma) + (1\|ID pa) | 63.83 |  |
| Egg protein ~ body weight + feed + egg size + (1\|tank) + (1\|ID ma) + (1\|ID pa) | 65.74 | 1.91 |
| Egg protein ~ body length + feed + egg size + (1\|tank) + (1\|ID ma) + (1\|ID pa) | 66.19 | 2.36 |

**Table S5.** Number of mature (stage IV) and immature (stages I – III) females by population, feed treatment, and number of families represented in the warm water treatment. Individual fish that had no population information available were excluded from the number of fish by population (*n* = 11).

|  |  | *n* | Immature | Mature |
| --- | --- | --- | --- | --- |
| **Population** |  |  |  |  |
| Neva |  | 346 | 302 | 44 |
| Oulu |  | 443 | 432 | 11 |
| **Feed treatment** |  |  |  |  |
| Control |  | 383 | 355 | 28 |
| Low-fat |  | 417 | 386 | 31 |
| **Families** |  | 84 | 84 | 23 |

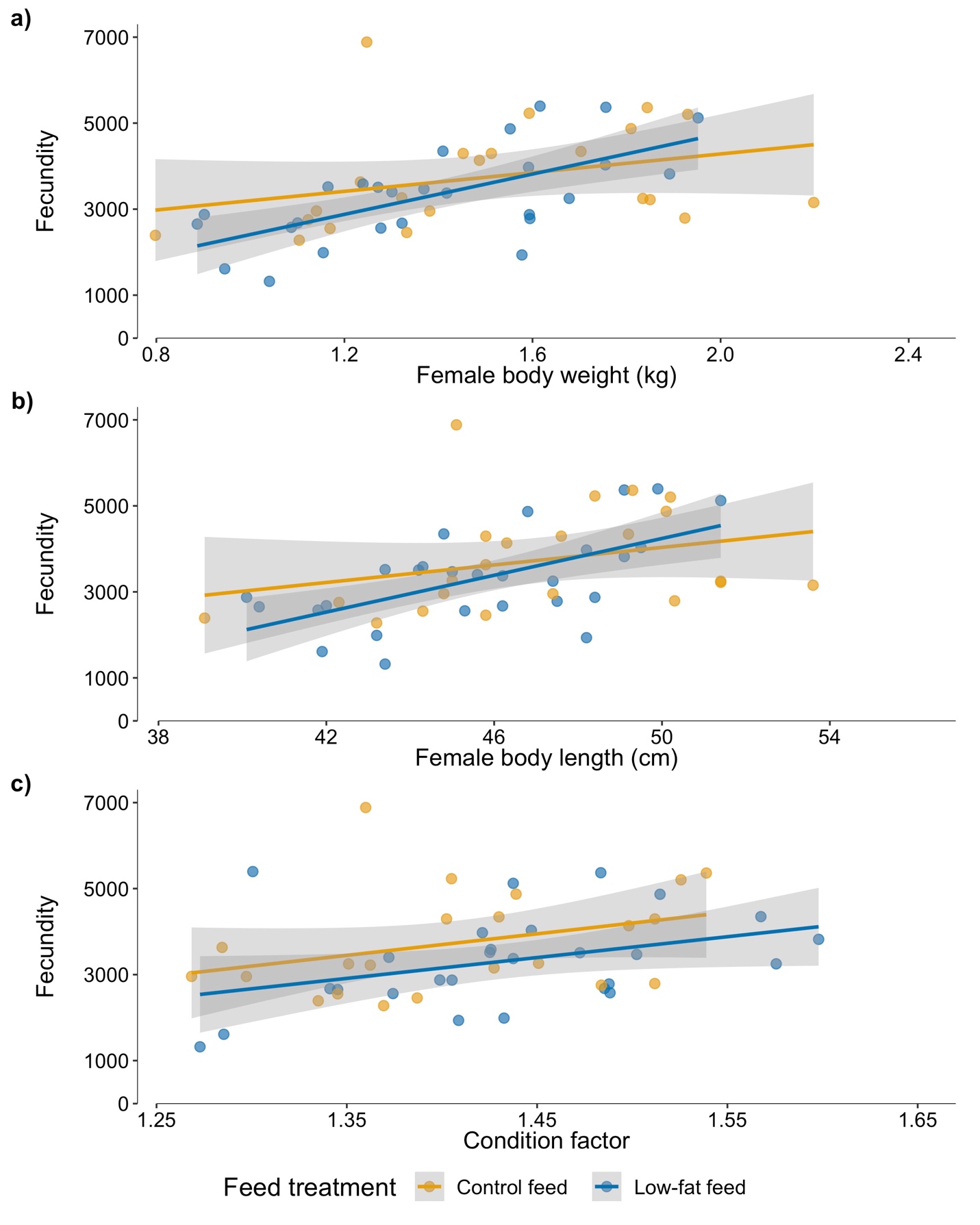

**Fig. S3.** Relationship between female a) wet weight, b) length and c) condition factor and fecundity (egg number) in the high energy and low energy feed treatments. The lines represent a linear regression and the grey area the 95% confidence interval.
